# MultiFRAGing: Rapid and Simultaneous Genotyping of Multiple Alleles in a Single Reaction

**DOI:** 10.1101/523837

**Authors:** Cassidy Petree, Gaurav K Varshney

**Affiliations:** Functional & Chemical Genomics Program, Oklahoma Medical Research Foundation, Oklahoma City, OK 73104

**Author notes:** Correspondence should be addressed to GKV.

## Abstract

The powerful and simple RNA-guided CRISPR/Cas9 technology is a versatile genome editing tool that has revolutionized targeted mutagenesis. CRISPR-based genome editing has enabled large-scale functional genetic studies through the generation of gene knockouts in a variety of model organisms including zebrafish. CRISPR/Cas9 can also be used to target multiple genes simultaneously. One of the challenges associated with applying this technique to zebrafish in a high-throughput manner is the absence of a cost-effective method by which to identify mutants. To address this, we optimized the high-throughput, high-resolution fluorescent PCR-based fragment analysis method to develop MultiFRAGing, a robust and cost-effective method for genotyping of multiple targets in a single reaction. Our approach can identify indels in 4 targets from a single reaction, which represents a four-fold increase in genotyping throughput. This method can be used by any laboratory with access to capillary electrophoresis based sequencing equipment.

## Introduction

Following completion of the human genome sequencing project, the identification of candidate disease genes has been the focus of much genetic research. With the development of less expensive sequencing technologies, these genes are being discovered at a rapid rate, however functional validation remains slow. Most of the knowledge of gene function has been elucidated from model organisms using gene knockout technology ^1^. The zebrafish (*Danio rerio*) has become a popular model organism for many reasons including high fecundity, optically transparent embryos and larvae, external development and the ease with which various types of genetic manipulation can be performed ^2^. Recent progress in the transformative, targeted, and simple RNA-guided CRISPR/Cas9-based genome editing method has expedited genetic manipulation in many systems, including zebrafish ^3–5^. These targeted mutagenesis methods are being used to generate knockouts for the purposes of developing disease models and understanding disease pathology in zebrafish. Many workflows are available for generating large numbers of gene knockouts ^6–10^, and while the generation of knockouts using CRISPR/Cas9 is relatively straight forward, the identification of mutations in a high-throughput, affordable manner remains a challenge. More than 70% of the indels generated by CRISPR/Cas9 are less than 10 bp long, making genotyping difficult ^6^. The most sensitive method used to identify indels involves amplification of the target region followed by cloning and sequencing; a process which is labor intensive and time consuming. A number of other methods such as high resolution melt analysis (HRMA), restriction fragment length polymorphism (RFLP) analysis, PAGE-based screening, and Surveyor assay all of which are suitable to identify mutations on a small scale ^11–14^. We and others previously adopted a high-throughput, high-resolution fragment analysis-based method to genotype CRISPR-induced alleles ^6^, ^15^, and later showed that it can also be used to determine guide RNA activities *in vivo* ^*^16^*^. The fragment analysis involves PCR-generation of fluorescently labeled fragments, and subsequent separation by size using capillary electrophoresis; software determines the relative size of each fluorescently labeled fragment by comparison with a size standard to generate the genotype of each ^17^. As the throughput of CRISPR/Cas9- mediated mutagenesis increases, researchers are able to target multiple genes simultaneously; this necessitates the development of a multiplex genotyping method to reduce both cost and labor. Here, we developed MultiFRAGing, a multiplexing fragment analysis pipeline that could genotype up to four targets in a single reaction, increasing the throughput by 3-4 fold while significantly reducing cost.

## Methods

### Zebrafish care and husbandry

All zebrafish experiments were carried out in compliance with the National Institutes of Health guidelines for animal handling and research under Oklahoma Medical Research Foundation (OMRF) Institutional Animal Care and Use Committee (IACUC) approved protocol 17-01. Wildtype (WT) zebrafish strain TAB-5 was used for all experiments. Zebrafish embryos were maintained in E3 embryo medium with 0.00002% methylene blue and raised at 28°C.

### Generation of mutant lines using CRISPR/Cas9 in zebrafish

The guide RNAs (sgRNAs), Cas9 mRNA and microinjections were carried out as described earlier ^6, 7^. Injected eggs were raised to the adulthood to generate founder fish. Six to eight founder fish were outcrossed with the wild type fish to generate heterozygous progenies (F_1_). Progenies from founders carrying mutations were raised to the adulthood to generate the F_1_ generation, adults of which were genotyped using fragment analysis as previously described ^6, 7, 17^ with the modifications described below.

### Multiplex PCR and Fragment Analysis

Primers were designed to amplify amplicons 200-300 bp in length and keep the target site in the middle. To generate fluorescently labeled fragments, we used M13F, T3 or SP6 sequences to tail gene-specific forward primers. A third fluorescently labeled primer, M13F-FAM, T3-TAMRA or SP6-HEX primer was designed to use together with gene-specific primers. This strategy allowed us to avoid the cost associated with the fluorescent labeling of individual primers. In order to avoid stutter peaks in genotyping, we added a 7-nucleotide long tag at the end of the gene-specific reverse primer. All three primers were mixed together as follows:

- 5µL 100 µM Fluorescent Primer
- 3µL 100 µM Forward Primer
- 5µL 100 µM Reverse Primer
- 487 µL water

PCR reactions were set-up in 20 µl final volume as follows:

- 2µL 10x Buffer
- 0.6µL MgCl2
- 0.4µl dNTP Mix
- 1.2µL Primer mix (for each)
- 0.16µl Platinum Taq Polymerase
- 2µL DNA Template
- To 20µL with Water

PCR was performed using the following conditions:

5 min denaturation at 94°C followed by 40 cycles of: 94°C for 30 sec, 57°C for 30 sec, and 72°C for 30 sec; and 5 min final extension at 72°C. Successful PCR amplification was confirmed using electrophoresis using 2% agarose gels, and amplified fragments were separated by capillary electrophoresis on a Genetic Analyzer. The ROX400 size standard (Thermo Fisher Scientific) was used as an internal size marker: ROX400 was diluted 1:100 in HiDi formamide and 7.5 µl of diluted mix was added to 2.5 µl of pooled PCR product. Samples were mixed and then denatured at 95°C for 5 minutes before separation on the Genetic Analyzer. Allele sizes were determined using GeneMapper software (Thermo Fisher Scientific).

### Verification of Alleles by Sequencing

To determine the nature of indels, PCR products from the fragment analysis reaction were directly sub-cloned into a pCR-TOPO4 vector. Plasmid was extracted using a Zymo Plasmid Miniprep kit (Zymo Research), and 100 ng DNA from each of 4 clones were subjected to sequencing using a BigDye Terminator Cycle Sequencing kit (Applied Biosystems). The resulting DNA fragments were purified and sequenced using a Capillary Electrophoresis Sequencer 3500Xl (Applied Biosystems), and sequences were aligned with wild type using SnapGene software (GSL Biotech LLC).

## Results

Our aim was to establish a reliable multiplex method by which to identify indels from multiple targets in a single PCR reaction to save time and cost, and to increase genotyping efficiency (Figure 1). The fragment analysis workflow presented here involves labeling fragments with fluorescent dyes to allow multiple colors of fluorescent dyes to be detected in a single sample; one of the dye colors is used for a size standard. ABI genetic analyzers can accommodate at least 4 different fluorescent dyes. We used the DS-30 dye set with 6-FAM (blue), Hex (green), NED (yellow), and Rox (Red). Rox was used as the labeled size standard, leaving the three others available. (If more colors are needed, the DS-33 dye set - which contains a fifth color - can be used.) As described in the methods section, gene-specific forward primers tailed with an adapter sequence (which lacks similarity to the genome) are used; in this case M13F, T3 and SP6 sequences. The same sequences were used for dye-labeling. We replaced the NED dye with TAMRA because it was readily available and inexpensive to synthesize.

**Figure 1:**
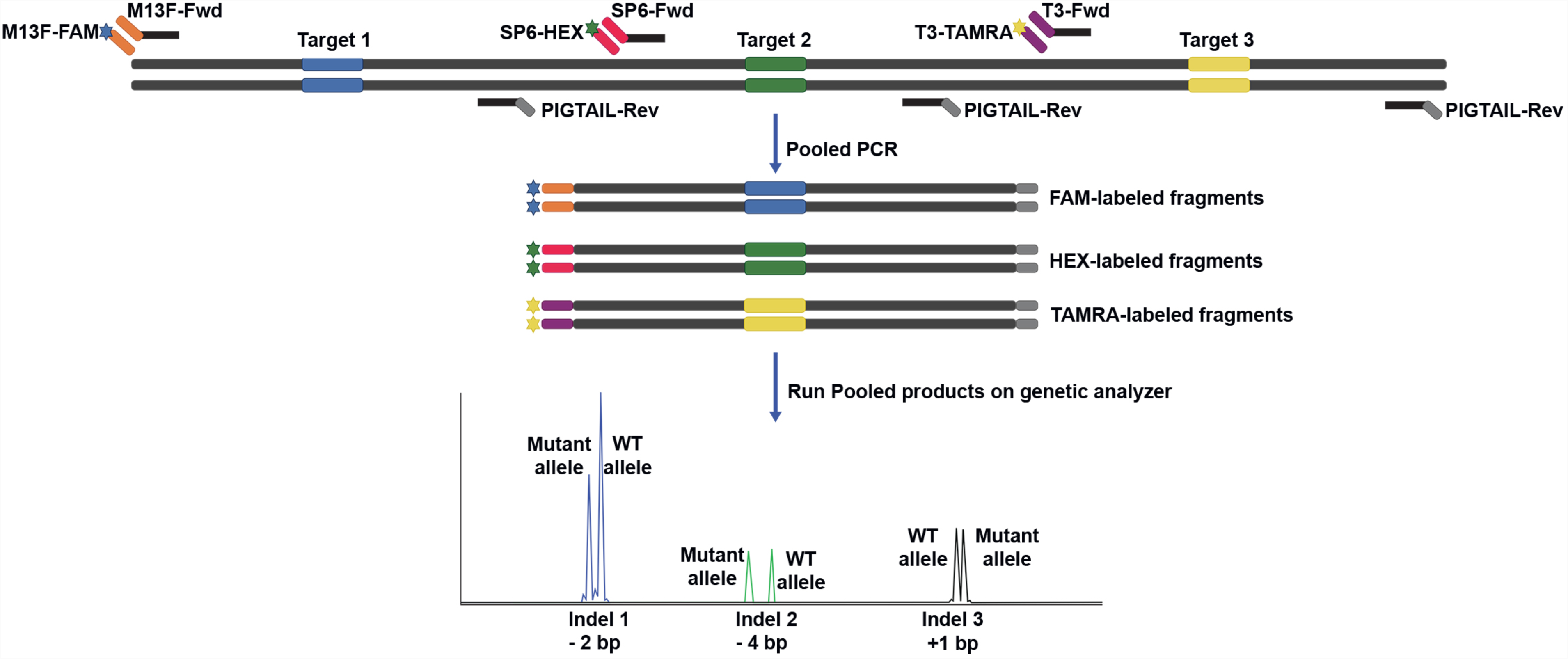
Overview of MultiFRAGing method. **A.** Primer design strategy for multiple targets. Gene specific primers are designed to generate 200-300bp long fragments. The gene-specific forward primers are attached with an adapter sequence (M13Fwd, SP6, and T3), and the reverse primer contains a short PIGTAIL sequence to suppress stutter. A third primer with same adapter sequence attached to a fluorescent dye (FAM, HEX and TAMRA) is used to generated fluorescently labeled fragments. **B.** Multiple fragments are generated in a single PCR reaction. These fragments can either be tagged with same dye but different size or tagged with different dyes. **C.** Pooled PCR products are then mixed with a size standard to run on a genetic analyzer. Fragments sizes are plotted and based on expected wild type fragment size, indel size can be measured.

It has been shown that the majority of indels induced by CRISPR/Cas9 are less than 10 bp long. This makes it possible to design specific primers to generate fragments of different sizes (within a 200-300bp range), thereby allowing us to vary both fragment length and dye color (Table1). Based on these two parameters we tested following combination of fragments: fragments labelled with two or three colors, fragments of two or three sizes, and fragments with different sizes and color together. To establish this method we used a mutant line carrying mutations at five distinct sites in four different genes (*Dfnb31a T1, Dfnb31bT1, Grhl2a T1, and Grhl2b T1*). First, we genotyped these alleles separately and confirmed the five independent alleles listed in Table 2. These alleles were then used to develop the multiplex method retrospectively. We amplified up to four targets in a single reaction using different DNA polymerases (Platinum Taq, AmpliTaq Gold), including one specialized for multiplex PCR (NEB Multiplex PCR 5X Master Mix, and Phusion Multiplex PCR Master Mix). Surprisingly, standard polymerases were more effective than multiplex PCR master mix, and Platinum Taq performed slightly better than AmpliTaq Gold. We successfully amplified 3 targets simultaneously (Supplementary Figure 1). The PCR products were combined with a size standard, and run on capillary electrophoresis to identify indels. As shown in Figure 2, we were able to identify indels using different combinations from the same PCR reaction. Therefore, this strategy of amplifying three targets simultaneously can increase the genotyping throughput three-fold.

**Table 1:**
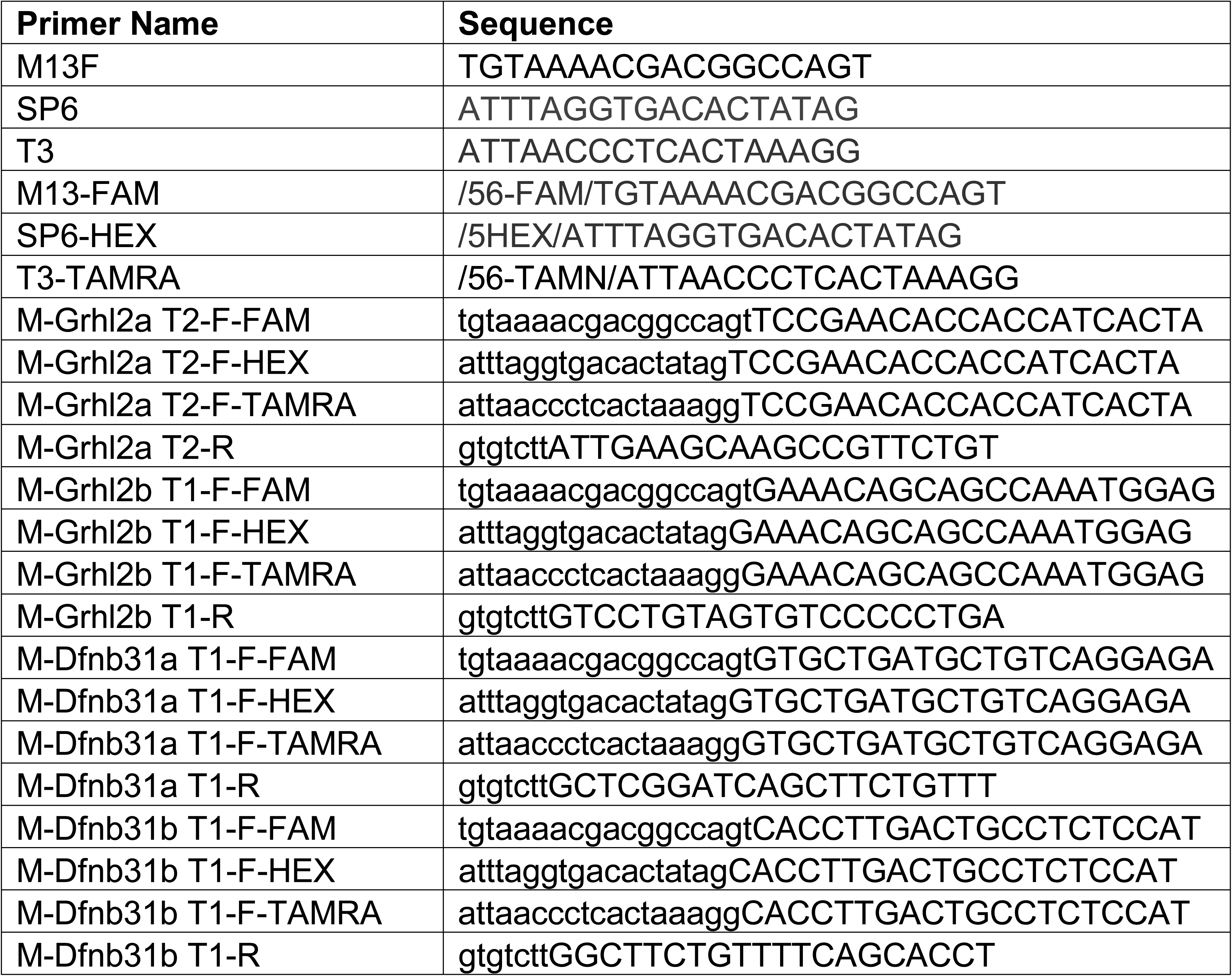
Primer sequences used in this study.

**Table 2.**
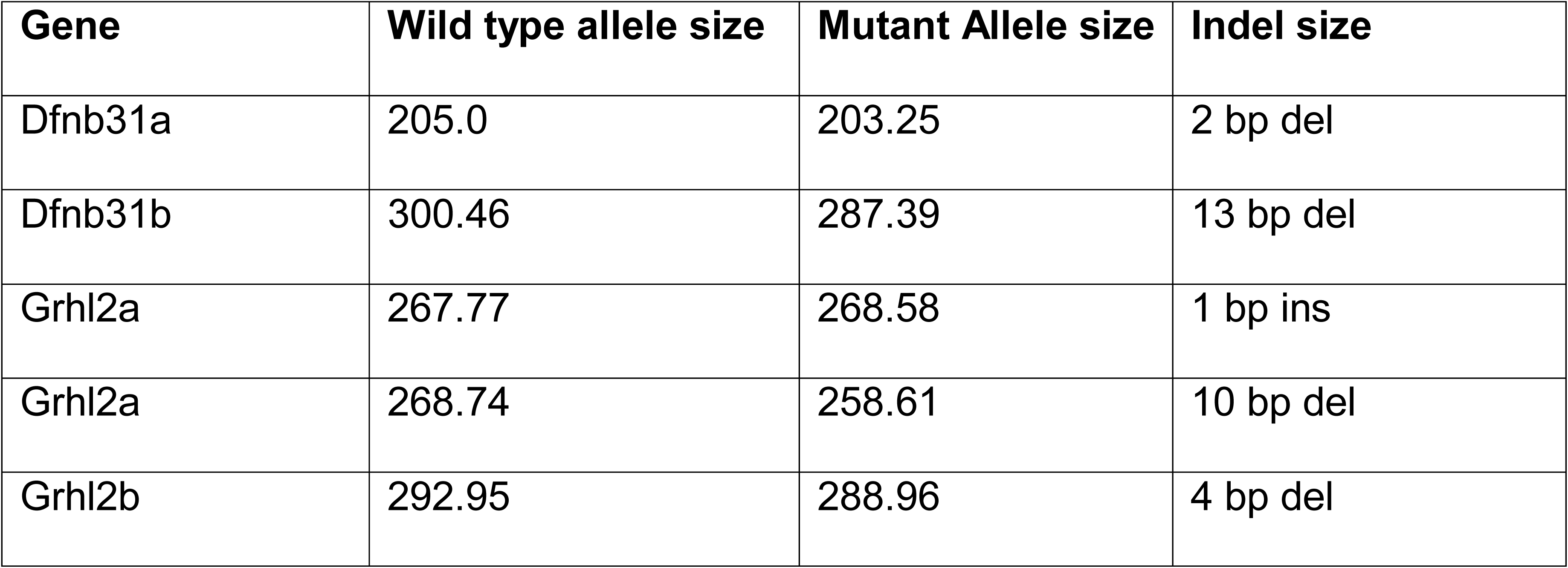
Summary of wild type, mutant allele, and predicted indel sizes used in this study.

| **Gene** | **Wild type allele size** | **Mutant Allele size** | **Indel size** |
| --- | --- | --- | --- |
| Dfnb31a | 205.0 | 203.25 | 2 bp del |
| Dfnb31b | 300.46 | 287.39 | 13 bp del |
| Grhl2a | 267.77 | 268.58 | 1 bp ins |
| Grhl2a | 268.74 | 258.61 | 10 bp del |
| Grhl2b | 292.95 | 288.96 | 4 bp del |

**Figure 2:**
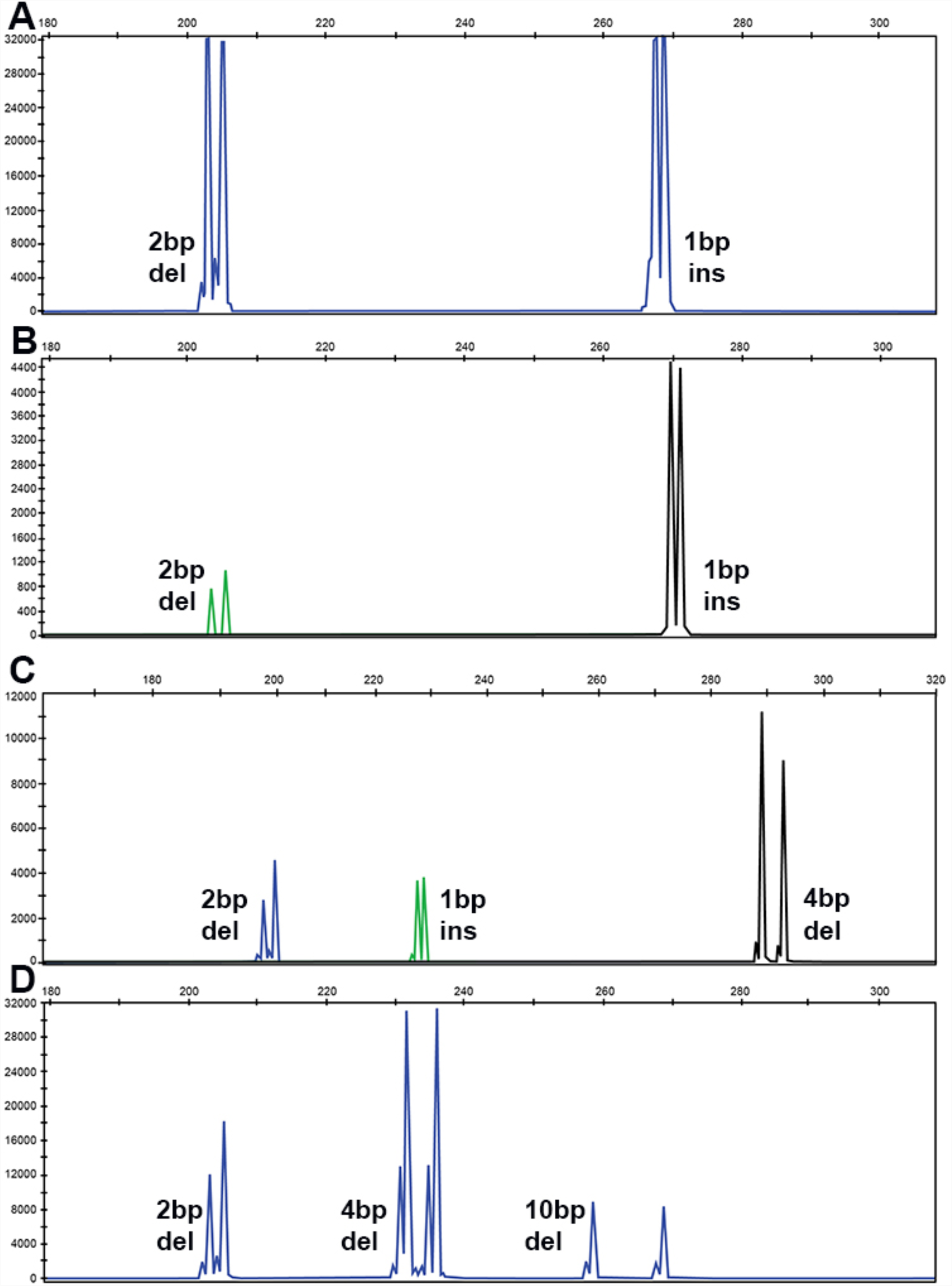
Fragment Analysis PCR plots from a pool of samples amplified together. **A).** Two fragments of different size labeled with FAM dye showing a 2bp deletion, and 1bp insertion. **B)** Two fragments are separated based on different dyes. **C)** Three fragments are separated by different dyes and 2bp deletion, 1bp insertion, and 4 bp deletion was detected **D)** Three fragments with same color and different sizes amplified together and analyzed as a pool in a single reaction.

Finally, we wanted to test whether individually amplified targets with different colors and/or sizes could be pooled together to allow indels to be identified in a single reaction. To test this approach, 2 µl from 3 or 4 different PCR products were pooled together, and 2.5 µl pooled product was sequenced from a single well. As shown in Figure 3, all alleles can be successfully identified, indicating that it is possible to pool multiple PCR products to increase the genotyping throughput up to four-fold and save the cost of consumables time, and labor. All alleles identified by fragment analysis were confirmed by the sequencing, and the sequencing data was in agreement with the fragment analysis data confirming the sensitivity of this method (Figure 4).

**Figure 3.**
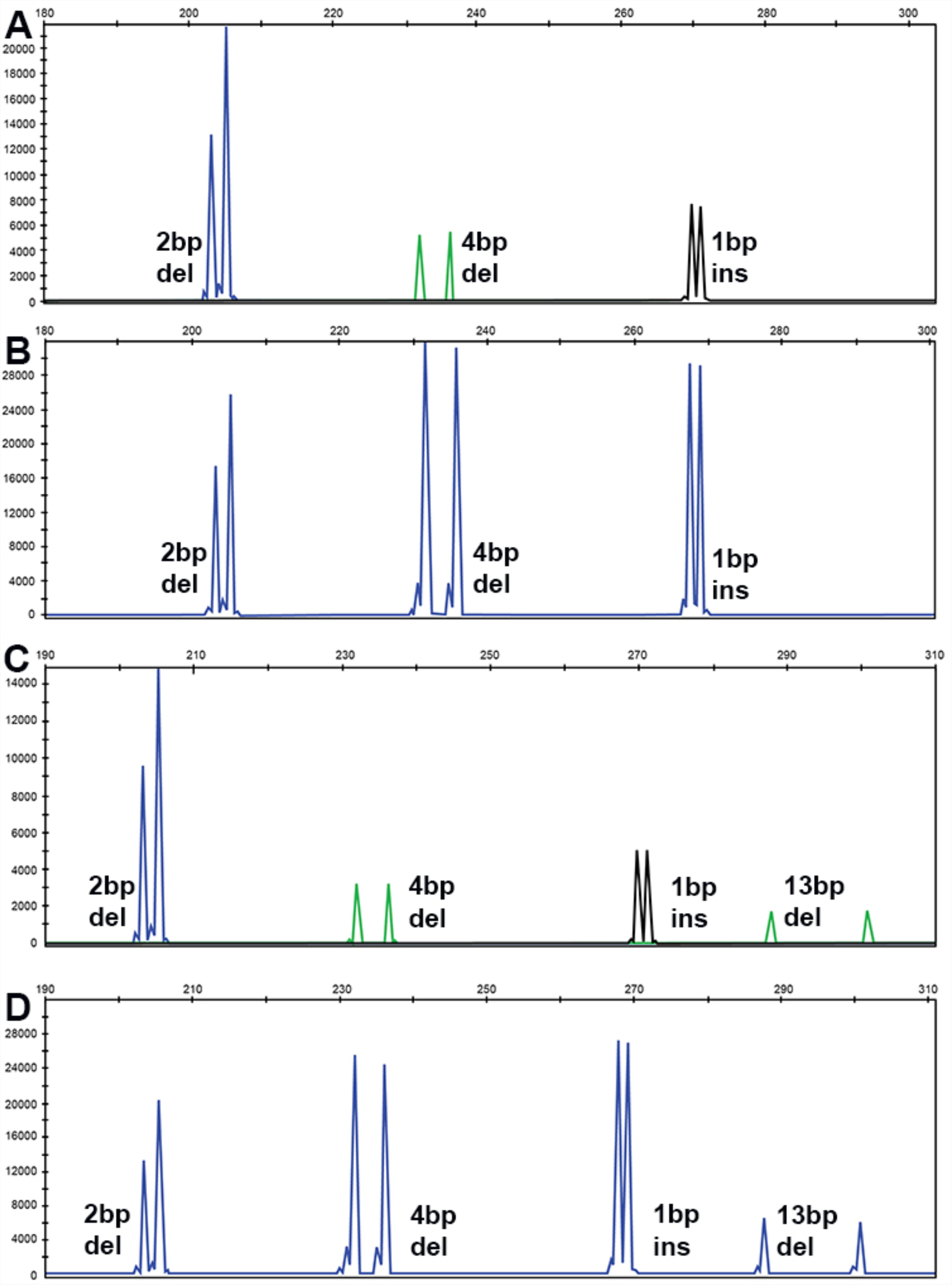
Fragment Analysis PCR plots from fragments generated by PCR individually and pooled together for genetic analyzer. **A**) Three fragments of different sizes but different dye colors showing indels of 2 bp deletion, 4bp deletion and 1bp insertion. **B)** Three fragments of different sizes but same dye color **C)** Four fragments of different size and different dye color showing indels of 2 bp deletion, 4bp deletion,1bp insertion, and 13bp deletion. **D).** Four fragments of different size but same dye color are separated.

**Figure 4.**
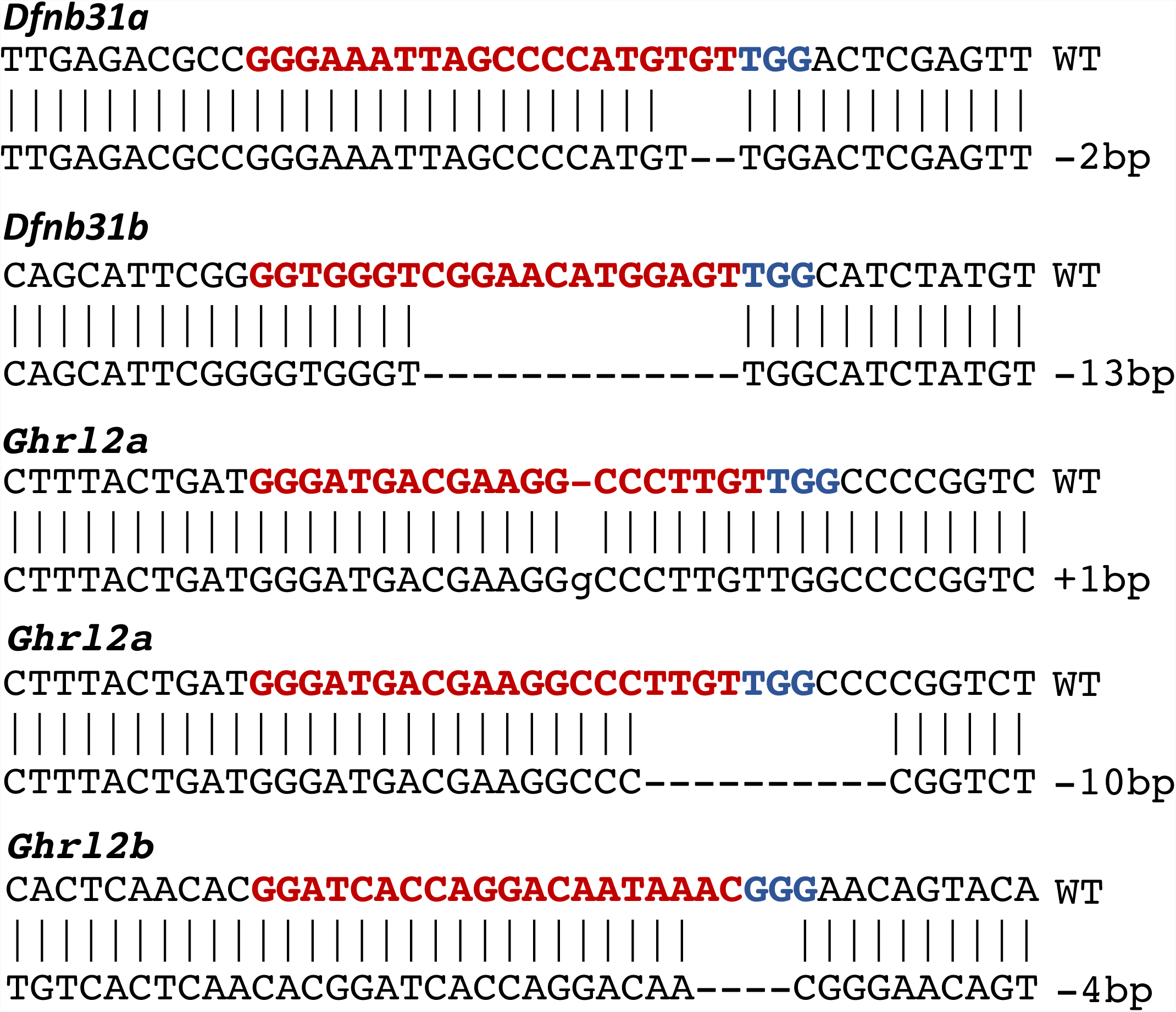
Validation of indels by Sanger Sequencing. Each indel that was identified by fragment analysis was sequenced by Sanger sequence to establish the correlation between fragment analysis and Sanger sequencing Data. All indels from fragment analysis showed similar indel size in Sanger sequencing.

In conclusion, our data shows that fragment analysis can be used to genotype multiple alleles simultaneously, making it a more robust and cost-effective genotyping method. Multiplexing can be done using two different approaches: amplify up to 3 targets in a single PCR reaction, and genotype the products in a single reaction, or amplify targets individually and pool the products for genotyping in a single reaction. We have demonstrated the use of multiplexing using zebrafish, but this method can also be adopted easily in other systems as well.

## Acknowledgements

We thank Rachel Smith for help with figures, Mackenzie Endebrock, Duane Goins and the staff of the Comparative Medicine department for animal husbandry work.

---

**Supplementary Figure 1:** Separation of PCR products from Pooled PCR on agarose electrophoresis. M: 100bp marker, Lane 1: Two different fragments with different sizes, same color. Lane 2: Two different fragments with different sizes, different color. Lane 3: Three different fragments with different sizes, same color. Lane 2: Three different *fragments* with different sizes, different color.

