## Supplementary figures and images for "MultiFRAGing: Rapid and Simultaneous Genotyping of Multiple Alleles in a Single Reaction"

### Supplementary File

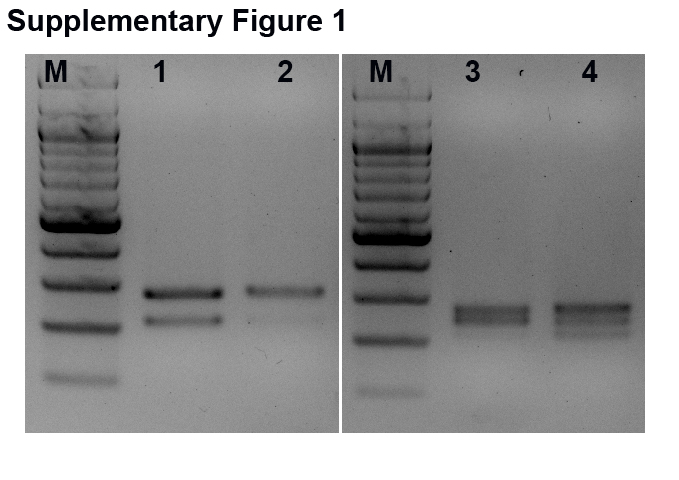
